# Speaker-normalized vowel representations in the human auditory cortex

**DOI:** 10.1101/397026

**Authors:** Matthias J. Sjerps, Neal P. Fox, Keith Johnson, Edward F. Chang

## Abstract

Humans identify speech sounds, the fundamental building blocks of spoken language, using the same cues, or acoustic dimensions, as those that differentiate the voices of different speakers. The correct interpretation of speech cues is hence uncertain, and requires normalizing to the specific speaker. Here we assess how the human brain uses speaker-related contextual information to constrain the processing of speech cues. Using high-density electrocorticography, we recorded local neural activity from the cortical surface of participants who were engaged in a speech sound identification task. The speech sounds were preceded by speech from different speakers whose voices differed along the same acoustic dimension that differentiated the target speech sounds (the first formant; the lowest resonance frequency of the vocal tract). We found that the same acoustic speech sound tokens were perceived differently, and evoked different neural responses in auditory cortex, when they were heard in the context of different speakers. Such normalization involved the rescaling of acoustic-phonetic representations of speech, demonstrating a form of recoding before the signal is mapped onto phonemes or higher level linguistic units. This process is the result of auditory cortex’ sensitivity to the contrast between the dominant frequencies in speech sounds and those in their just preceding context. These findings provide important insights into the mechanistic implementation of normalization in human listeners. Moreover, they provide the first direct evidence of speaker-normalized speech sound representations in human parabelt auditory cortex, highlighting its critical role in resolving variability in sensory signals.

## Introduction

A fundamental computational challenge faced by perceptual systems is the lack of a one-to-one mapping between highly variable sensory signals and the discrete, behaviorally relevant events they reflect[1,2]. A profound example of this problem exists in human speech perception, where the main cues to speech sound identity are the same as those to speaker identity[3–5].

For example, to distinguish a given speaker’s /*u*/ from his or her /*o*/ (distinguishing “boot” from “boat”), listeners rely heavily on the vowel’s first formant frequency (F1; the first vocal tract resonance) because it is lower for /u/ than for /o/[6]. However, people with long vocal tracts (typically tall male speakers) have overall lower resonance frequencies than those of speakers with shorter vocal tracts. Consequently, a tall person’s production of the word “boat” and a short person’s “boot” might be acoustically identical. Behavioral research has suggested that preceding context allows listeners to “tune-in” to the acoustic properties of a particular voice and *normalize* subsequent speech input[7–11]. The most well-known example of this effect is that a single acoustic token, ambiguous between /u/ and /o/, will be labelled as /o/ after a context sentence spoken by a tall-sounding person (low F1), but like /u/ after a context sentence spoken by a shorter-sounding person (high F1)[12].

The neurobiological foundations of context-based speaker normalization remain largely unknown. Neural activity in auditory cortex is sensitive to acoustic cues that are critical for both recognizing and discriminating phonemes[13–19] and different talkers[20–24]. For example, recent work has shown that speech sound representations in STG are closely related to the acoustic-phonetic features that define classes of speech sounds, like F1. Vowels with low F1 frequencies (e.g., /u/, /i/) can be distinguished from vowels with relatively higher F1 frequencies (e.g., /o/, /æ/) based on local activity on STG[25]. A critical question that arises, then, is whether the feature-based representations in auditory cortex are normalized (i.e., feature rescaling), or whether they continue to closely reflect the veridical acoustic properties of the input.

To investigate the influence of context on auditory cortex speech sound representations, we recorded cortical local field potentials with subdurally implanted high density electrode arrays that covered the broader peri-sylvian language region in human participants while they listened to and identified vowel sounds presented in the context of sentences spoken by two different voices[14,26]. We found direct evidence of speaker-normalized neural representations of vowel sounds in parabelt auditory cortex, including superior and middle temporal gyri. Normalization was observed in populations that were selective for acoustic-phonetic (i.e., pre-phonemic) properties of the speech signal. These effects were at least partly driven by the contrastive relation between the F1 range in the context sentences and F1 values in the target vowels. More generally, the results demonstrate the critical role of human auditory cortex in integrating incoming sounds with surrounding acoustic context.

## Results

We recorded neural activity directly from the cortical surface of five Spanish-speaking neurosurgical patients while they voluntarily participated in a speech sound identification task. They listened to Spanish sentences that ended in a (pseudoword) target, which they categorized as either “sufu” or “sofo” on each trial with a button press (Figure 1a, b). The sentence-final targets comprised a digitally synthesized six-step continuum morphing from an unambiguous *sufu* to an unambiguous *sofo*, with four intermediate tokens (*s?f?*, i.e., spanning a perceptually ambiguous range). On each trial, a pseudo-randomly selected target was preceded by a context sentence (*A veces se halla…*; “*At times she feels rather*…”). Two versions of this context sentence were synthesized, differing only in their mean F1 frequencies (Figure 1a, c; Figure S1), yielding two contexts that listeners perceive as consistent with two speakers: one with a long vocal tract (low F1; Speaker A) and one with a short vocal tract (high F1; Speaker B). Critically, F1 frequency is the primary acoustic dimension that distinguishes between the vowels /*u*/ and /*o*/ in natural speech (in both Spanish and English), as well as in our target continuum (Figure 1a and Figure S1)[6]. Similar materials have previously been shown to induce a reliable shift in the perception of an /u/ – /o/ continuum (a “normalization effect”) in healthy Spanish-, English-, and Dutch-listeners[8].

**Figure 1:**
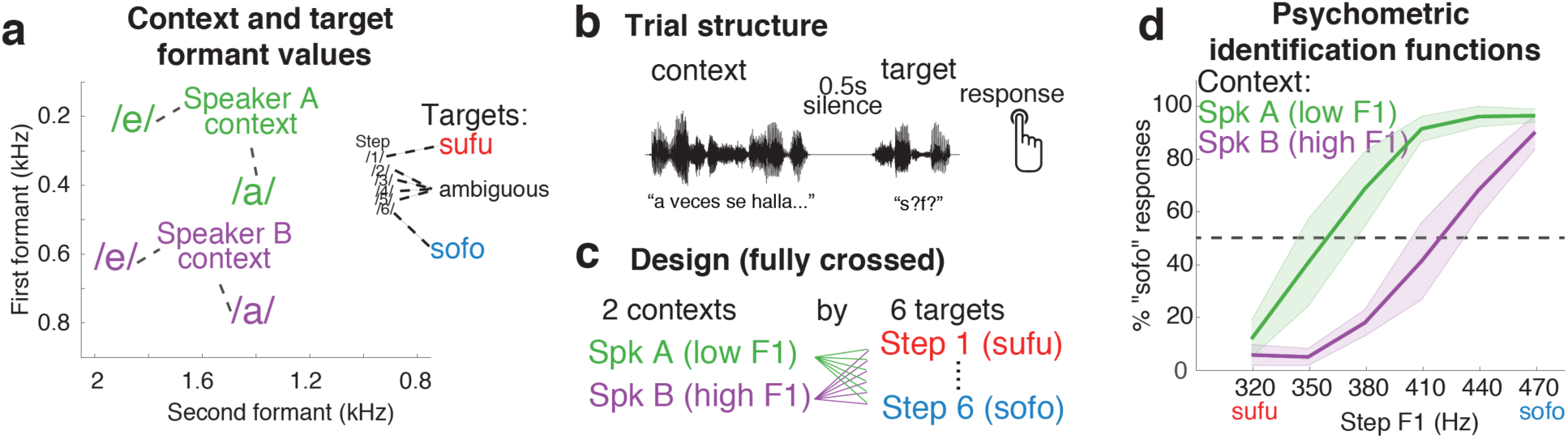
Listeners perceive speech sounds relative to their acoustic context. **a**) Target sounds were synthesized to create a 6 step continuum ranging from /sufu/ (step 1; low first formant [F1]) to /sofo/ (step 6; high F1). Context sentences were synthesized to sound like two different speakers: a speaker with a long vocal tract (low F1 range: Speaker A), and a speaker with a short vocal tract (high F1 range; Speaker B). Context sentences contained only the vowels /e/ and /a/, but not the target vowels /u/ and /o/. Following phonetic convention, the formant axes are reversed (higher values at the bottom/left sides of the panel). **b**) Context sentences preceded the target on each trial (separated by 0.5 seconds of silence), after which participants responded with a button press to indicate whether they heard “sufu” or “sofo”. **c**) All targets were presented after both speaker contexts. **d**) Listeners more often gave “sofo” responses to target sounds if the preceding context was spoken by Speaker A (low F1) than Speaker B (high F1).

As expected, participants’ perception of the target continuum was affected by the F1 range of the preceding sentence context (p < 0.002; Figure 1d). Specifically, participants were more likely to identify tokens as *sofo* (the vowel category corresponding to higher F1 values) after a low F1 voice (Speaker A) compared to the same target presented after a high F1 voice (Speaker B). Hence, listeners’ perceptual boundary between the /*u*/ and /*o*/ vowel categories shifted to more closely reflect the F1 range of the context speaker. Past work has interpreted this classical finding in light of the contrastive perceptual effects that are ubiquitous among sensory systems[27]: the F1 of a speech target will sound relatively higher (i.e., sound more like an /o/) after a low F1 context sentence than after a high F1 context. This results in a shift of the category boundary to lower F1 values.

### Human auditory cortex exhibits context-dependent speech sound representations

Two of the most influential hypotheses explaining the phenomenon of speaker normalization posit that: 1) contrast enhancing processes, operating at general auditory processing levels, change the representation of the input signal before it is mapped onto phonemes or higher level linguistic units[28–30]; 2) alternatively, it has been suggested that auditory processing of speech cues remains mostly faithful to the acoustics of the input signal, and normalization is a consequence of speaker-specific mapping of the veridical acoustics onto meaningful units (i.e., listeners have learned to associate an F1 of 400Hz to /u/-words for speakers, or vocal tract, that sound short, but to /o/-words for ones that sound taller)[9,31].

Past neurobiological work has demonstrated that neural populations in the parabelt auditory cortex are sensitive to acoustic-phonetic cues that distinguish classes of speech sounds, including vowels, and not to specific phonemes per se[25]. Hence, the primary goal of the current study was to examine whether the F1 range in preceding context sentences influence the representation of speech sounds in parabelt (nonprimary) auditory cortex in a normalizing way. We investigated whether the neural representation of vowel stimuli remains veridical (i.e., unaffected by context) or, alternatively, whether it becomes shifted towards the representation typical of /u/ in the context of a high F1 speaker, but towards /o/ in the context of a low F1 speaker. We first tested whether individual cortical sites that reliably differentiate between vowels (i.e., discriminate /u/ from /o/ in their neural response) would exhibit normalization effects. A secondary goal of the current study was to confirm that the response profile of those cortical populations that display normalization was indeed acoustic-phonetic (i.e., pre-phonemic) in nature. We therefore assessed populations’ responsiveness during the context sentences as well (context sentences did not contain the target phonemes /o/ and /u/ but did traverse the same acoustic F1 region).

To this end, we extracted the stimulus-aligned analytic amplitude of the high-gamma band (70-150 Hz) of the local field potential at each temporal lobe electrode (n = 406 across patients; this number is used for all Bonferroni corrections below) during each trial. High-gamma activity is a spatially- and temporally-resolved neural signal that has been shown to reliably encode phonetic properties of speech sounds[25,32,33], and is correlated with local neuronal spiking[34–36]. We used general linear regression models to identify cortical patches involved in the representation of context and/or target acoustics. Specifically, we examined the extent to which high-gamma activity at each electrode depended on stimulus conditions during presentation of the context sentences (context window) or during presentation of the target (target window; see supplemental materials). The fully specified encoding models included numerical variables for the target vowel F1 (Steps 1-6) and context F1 (high vs. low), as well as their interaction. In the following, we focused on “task-related electrodes”, defined as the subset of temporal lobe electrodes for which a significant portion of the variance was explained by the full model, either during the target window or the during context window (p < 0.05; uncorrected, n = 98; see Figure S2).

Among the task-related electrodes, some displayed selectivity to target vowel F1 (Figure 2a). Consistent with previous reports of auditory cortex tuning for vowels[25] we observed that different subsets of electrodes displayed a preference for either “sufu” or “sofo” targets (color coded in Figure 2a). Figure 2b and Figure 2c (middle panel) display the response profile for one example electrode that had a “sofo” preference (e1; *p* = 6.8*10^-19^). Importantly, in addition to an overall selectivity to the target sound F1, the activation level of this electrode was modulated by the F1 range of the *preceding* context (Figure 2b & bottom panel of Figure 2c; *p* = 5.8*10^-6^). This demonstrates that the responsiveness of a neural population that is sensitive to bottom-up acoustic cues is also affected by the distribution of that cue in preceding context. The direction of this influence is analogous to the behavioral normalization effect.

**Figure 2:**
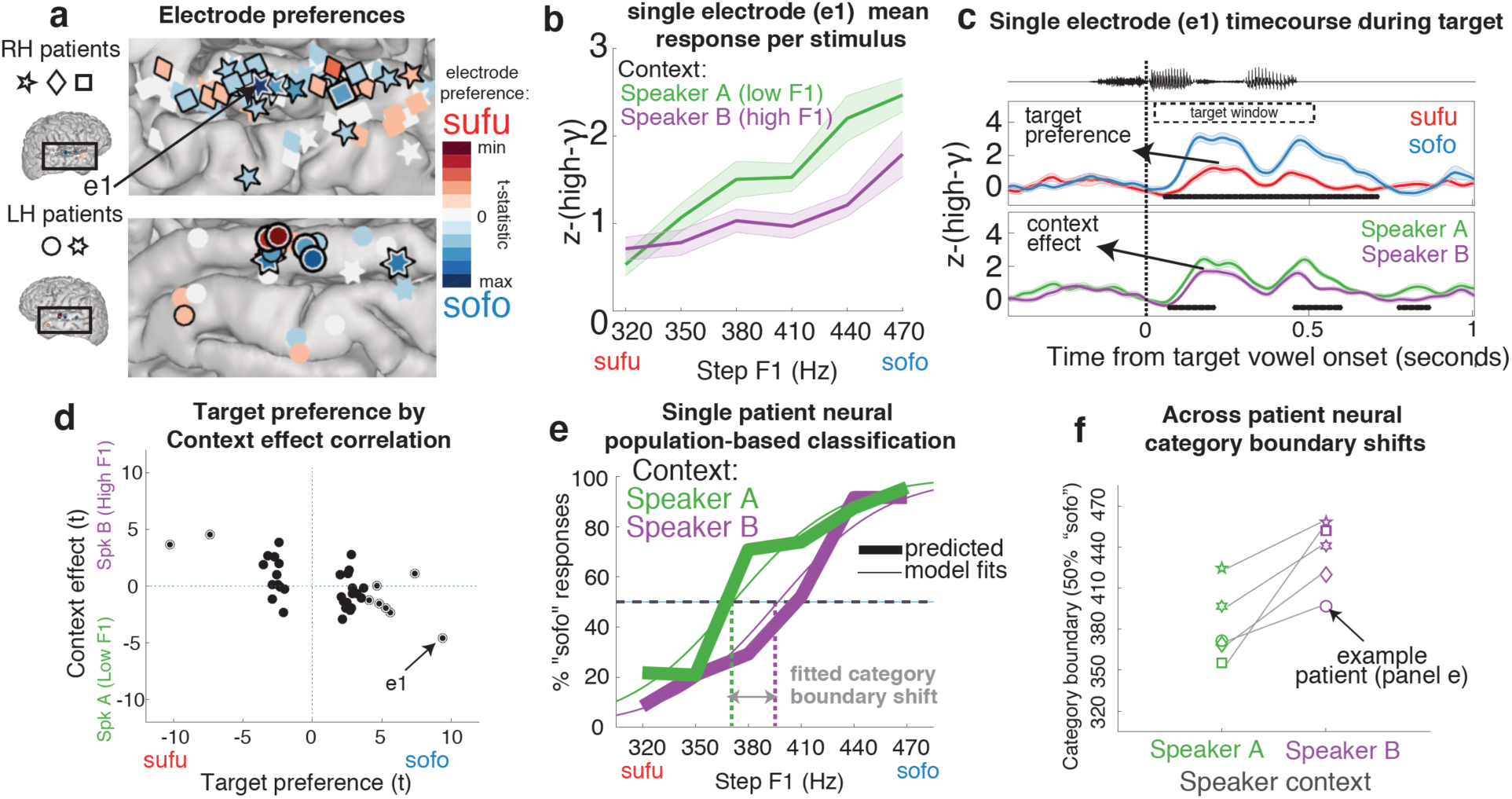
The neural response to bottom-up acoustic input is modulated by preceding context. **a**) Target vowel preferences and locations (plotted on an MNI brain) for electrodes from all patients (3 with right hemisphere [RH] and 2 with left hemisphere [LH] grid implants). Only those temporal lobe electrodes where the full omnibus model was significant during the context and/or the target window (F-test; p<0.05) are displayed. Strong target F1 selectivity is relatively uncommon: electrodes with a black-and-white outline are significant at Bonferroni corrected p<0.05 (n = 9, out of 406 temporal lobe electrodes); a single black outline indicates significance at only p<0.05, uncorrected (n = 28). Activity from the indicated electrode (e1) is shown in b and c. **b**). Example of normalization in a single electrode (e1; z-scored high-gamma [high-γ] response averaged across the target window [target window marked in c]; +-1SE). **c**) Activity from e1 across time, separating the endpoint targets (top panel) or the contexts (bottom panel). The electrode responds more strongly to /o/ stimuli than /u/ stimuli, but also responds more strongly overall after Speaker A (low F1). This effect is analogous to the behavioral normalization (Fig. 1d). Black bars at the bottom of the panels indicate significant time-clusters (cluster-based permutation test of significance). **d**) Among all electrodes with significant target sound selectivity (n = 37 [9 + 28]), a relation exists between the by-electrode context effect and target preference. Both are expressed as a signed t-value, demonstrating that the size and direction of the target preferences predicts the size and direction of the context effects. As per a, symbol style reflects level of significance (solid back = p<0.05 uncorrected; black-and-white = p<0.05 Bonferroni corrected). **e)** An LDA classifier was trained on the distributed neural responses elicited by the “sufu” and “sofo” stimuli using all endpoint selective electrodes of a patient. This model was then used to predict classes for (held-out) endpoint data and for the ambiguous steps. Proportions of neurally-based “sofo” predicted trials (thick lines) display a relative shift between the two context conditions (data from one example patient). Regression lines were fitted to these data for each participant separately to estimate 50% category boundaries per condition for panel f (thin lines). **f**) The neural classification functions display a shift in category boundaries between context conditions for all patients individually. Symbols denote individual participants.

To quantify this normalization effect across all electrodes that display selectivity to target acoustics, we calculated the correlation between electrodes’ *target preference* (numerically defined as the glm-based signed t-statistic of the target F1 factor during the target window) and their *context effect* (defined as the t-statistic of the context F1 factor during the target window). We found a correlation between electrodes’ target preferences and context effects (Figure 2d). Crucially, this strong relationship had a negative slope, such that electrodes that had high-F1 target preferences (sofo > sufu) had stronger responses to targets after low F1 context sentences (low F1 context > high F1 context; *r* = -0.65; *p* = 1.3*10^-6^). Importantly, this demonstrates that the relationship between context response and target response reflects an encoding of the contrast between the formant properties of each, recapitulating the normalization pattern observed in the behavioral responses (Fig 1d).

### Normalization of distributed vowel representations in all participants

Figure 2d demonstrates that local populations in auditory cortex that are selective to target vowel F1 exhibit normalization. However, only a few electrodes (n = 9, out of 406 temporal lobe electrodes) displayed very strong target vowel selectivity (significance at Bonferroni-corrected p <0.05), while the majority of target F1 selective electrodes displayed only moderate selectivity and context effects. Moreover, not all participants had such highly target F1 selective electrodes (see Table S2). The relative sparseness of strong selectivity is not surprising given that the target vowel synthesis involved only small F1 frequency differences (∼30Hz) per step, with the endpoints being separated by only 150Hz (which is, however, a prototypical F1 distance between /u/ and /o/[8]). However, past work has demonstrated that even small acoustic differences among speech sounds are robustly encoded by distributed patterns of neural activity across auditory cortex[14,37]. In order to determine whether distributed neural representations of vowels reliably display normalization across all participants, we trained a multivariate pattern classifier model (Linear Discriminant Analysis, LDA) on the spatiotemporal neural response patterns of each participant. Models were trained to discriminate between the endpoint stimuli (i.e., trained on the neural responses to steps 1 vs. 6, irrespective of context) using all task-related electrodes for that participant. These models were then used to predict labels for held-out data of both the endpoints and the ambiguous steps. For all participants, classification of held-out endpoint trials was significantly better than chance (Figure S3b). To assess the influence of target F1 and context F1 on the classifier output, a logistic generalized linear mixed model was then fit to the proportion of predicted “sofo” responses across all participants.

Figure 2e displays the proportion of “sofo” labels predicted for all stimuli by the LDA classifier based on the neural data of one example participant (thick lines). Importantly, a shift is observed in the point of crossing of the category boundary. Regression functions fitted to these data (thin lines) allowed us to estimate the size and direction of the context-driven neural boundary (50% crossover point) shift per participant. For all participants, the neural vowel boundaries were found to be context dependent (Figure 2f; see online methods and Supplementary Figure S3 for further detail). The combined regression analysis demonstrated that, across participants, population neural activity in the temporal lobe was modulated both by the acoustic properties of the target vowel (p = 1.2*10^-7^) and by the preceding context (p = 4.2*10^-8^). This effect was not dependent on the approach to exclusively train on endpoint data (Figure S5). Moreover, this effect was not observed for task-related electrodes outside of the temporal lobe during the target window (see S4; non-temporal electrodes were mostly located on sensorimotor cortex and the inferior frontal gyrus).

Importantly, and in analogy to the behavioral results, the neural classification functions demonstrate that the influence of the context sentences consistently affected target vowel representations in a normalizing direction: the neural response of a target vowel with a given F1 is more like that of /o/ (high F1), after a low F1 context (Speaker A) than after a high F1 context (Speaker B; see Supplementary Figure S3 and S4b for more detail).

### Normalization as sensitivity to contrast in acoustic-phonetic features

It has been suggested that a major organizing principle of human parabelt auditory cortex concerns the acoustic phonetic features that define classes of speech sounds, and not phonemes (or even higher level linguistic representations) per se[13,25,38]. Here we demonstrated normalization in these representations. However, auditory cortex processing is diverse and may contain regions that are in fact selective for (more abstract) phonemes. For example, auditory cortex has been found to display properties that are typically associated with abstract sound categories such as categorical perception, too[14]. Hence, we next assessed whether the normalization effects observed here involved a rescaling in patches of cortex that display sensitivity to acoustic-phonetic features (i.e., relating to more general F1 characteristics) or, instead, only in those patches that may be selective for discrete phonemes (or the target words as a whole). Because the context sentence did not contain the target vowels /u/ or /o/, but did traverse the same general F1 range, assessing electrodes’ responses during the context window could inform us about the nature of their preferences.

To this end we again relied on the glm-based t-statistics of all target selective temporal lobe electrodes (n = 37; as per Figure 2d). Among these electrodes, however, we examined the relationship between their preferences for context F1 during the context window and for target F1 during the target window. Figure 3a displays context and target preferences on the cortex of a single example patient. Among the electrodes that displayed target F1 selectivity, some also displayed selectivity for the context F1 during the context window (indicated with a black-and-white outline). Figure 3b displays the activation profile of one example electrode (e2). Importantly, e2 responds more strongly to low F1 targets during the target window (*sufu* preferent: p = 2.4*10^-21^) but also to low F1 contexts during the context window (Speaker A preferent: p= 1.7*10^-14^). This demonstrates that this neural population responded more strongly to low F1 acoustic stimuli in general and is not exclusively selective for a discrete phoneme category. Importantly, e2 also displayed normalization, as its activity was affected by context F1 during the *target* window (p = 2.7*10^-4^), and the direction of that context effect was consistent with contrastive normalization (*cf*. Fig. 2d).

**Figure 3:**
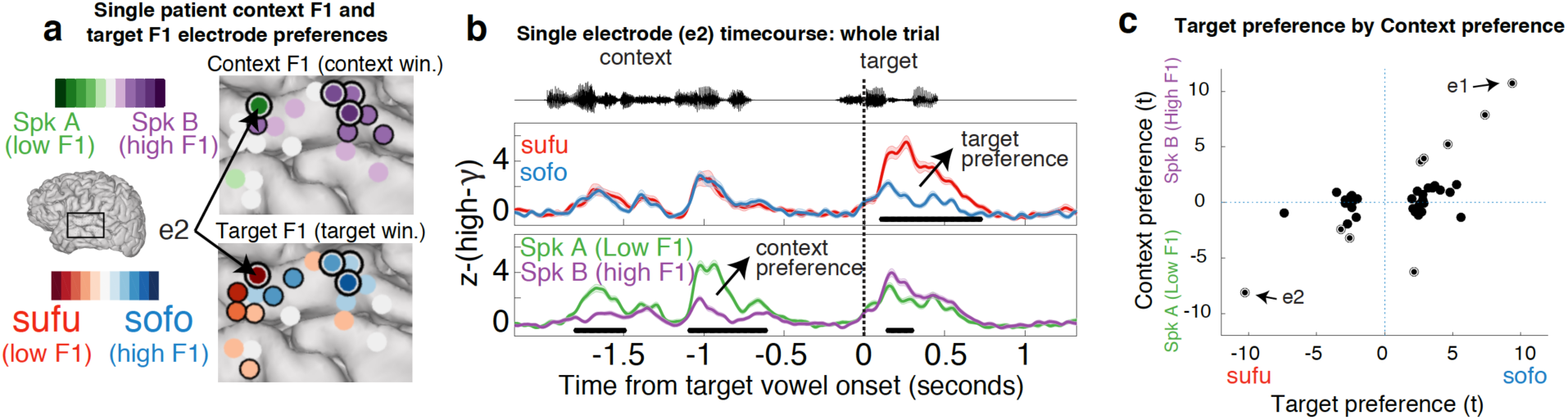
Sensitivity to contrast in acoustic-phonetic features. **a**) Electrode preferences for both target F1 (during target window) and context F1 (during context window) from a single example patient. Some populations display both target F1 selectivity and context F1 selectivity (marked with a black-and-white outline), indicating a preference for higher or lower F1 frequency ranges. Others are only selective for target F1 or context F1 (marked with a single black outline in their respective panels). Significance assessed at p<0.05 uncorrected. **b**) Mean (+/-1 SE) high-gamma activity at an example electrode (e2) from the example patient in panel a (conditions split as described in Fig 2c). Activity is displayed for a time window encompassing the full trial duration (both precursor sentence and target word). Black bars represent significant time points (p < 0.05; cluster-based permutation). **c**) A relation exists between the by-electrode context preference and target preference: electrodes that display a preference for either high or low target F1 typically also display a preference for the same F1 range during the context.

Extending this finding to the population of electrodes, we found a significant positive correlation across all target-selective temporal lobe electrodes between an electrode’s target preference and its context preference (*r* = 0.64; *p* = 1.42*10^-5^; Figure 3c). Hence, neural populations that are selective for target F1 in fact often displayed a more general preference to specific F1 frequency ranges. Moreover, when restricting the test of normalization (assessed as the correlation between target preferences and the context effect, as per Figure 2d) to those electrodes that displayed significant selectivity for both target F1 and context F1, normalization was again found (Figure S6). These findings confirm that normalization affects acoustic-phonetic (i.e., pre-phonemic) representations of speech sounds in parabelt auditory cortex.

## Discussion

A critical challenge for human speech perception is the fact that different speakers produce the same speech sounds differently[1,3]. That is, speakers display different effective ranges with respect to their most informative speech cues (here, formants). We investigated the neural underpinnings of the behavioral finding that listeners rely on speaker-specific information to constrain phonetic processing. First, we observed behavioral normalization effects, replicating previous findings[7,8,10,11]. More importantly, analogous speaker-normalized representations of vowels were found in parabelt auditory cortex processing. These normalized representations were observed broadly across parabelt auditory cortex and were observed for all participants individually. Normalization was found to involve a context dependent change in the response strength of cortical populations that are selective for acoustic-phonetic features. These findings demonstrate that normalization is a highly robust phenomenon that results in a rescaling of representations that precede the mapping onto phonemes or higher level linguistic units.

Recent research has demonstrated that auditory cortex responds to the acoustic cues that are critical for both recognizing and discriminating phonemes[13–19] and different talkers[20–24] by means of different patterns of activation[37]. However, since cues that are critical for speaker and speech sound identification are conflated in the acoustic signal, these findings could be consistent either with a cortical representation of veridical acoustic properties (e.g., reflecting the absolute F1 of a stimulus) or of context-dependent perceptual properties (e.g., its relative – or normalized – F1). Here, we were able to directly address the interaction between speaker and speech sound representations by presenting them to listeners at separate points in time and leveraging their immediate integration in auditory cortex processing. This approach demonstrated that rapid and broadly distributed normalization, or rescaling, is a basic principle of auditory cortex’s encoding of speech sounds.

In behavioral research on normalization, the effect has often been discussed in relation to its contrastive nature. Indeed, we observed that a low F1 context led to more high F1 target percepts, which is consistent with an increase of perceptual contrast. Similar contrast enhancing operations have been widely documented in human and animal processing of various (nonspeech) acoustic stimuli[39–42], involving phenomena such as adaptive gain control[41] or stimulus specific adaptation[40,42]. An intuitive mechanism for the implementation of contrast enhancement involves sensory adaptation. This could be based on neuronal fatigue. When a neuron, or neuronal population, responds strongly to a masker stimulus, its response during a subsequent probe is often attenuated when the frequency of the probe falls within the neurons’ excitatory receptive field[43,44]. But in addition to such local forms of adaptation, and possibly even more relevant for the effects observed here, adaptation also arises through (inhibitory) interactions between separate populations of neurons (which may have partly non-overlapping receptive fields)[39,41]. In the present study, spectral differences between the two context sentences and those between the endpoint target vowels were similar (see Figure S1). Adaptation may, hence, play a role in the type of normalization observed here. Indeed, we observed a number of populations for which a strong preference for one of the context sentences during the context window was associated with a decreased response during the target window (i.e., the normalization effect; Figure 3). Given the general nature of adaptation effects a relevant observation from the behavioral literature is the fact that various non-speech context sounds (e.g., broadband noise and musical tones) have also been observed to affect the perception of speech sounds in a way that is at least qualitatively similar to those observed here[28,29,45]. This finding suggests that normalization effects may not be speech specific, and may, at least partly, be explained by adaptation effects.

An interesting additional question concerns the main locus of emergence of normalization. Broadly speaking, normalization could be inherited from primary auditory or subcortical processes (from which we were unable to record); it may largely emerge within parabelt auditory cortex processing itself; or it could be driven by top-down influences from regions outside of the auditory cortex. In our study, context and target sounds were separated in time by a 500ms silent interval. It has been suggested that adaptation effects over such longer latencies become especially dominant at cortical levels of processing[46,47]. Furthermore, behavioral experiments have demonstrated robust normalization effects with contralateral presentation of context and target sounds[28,45]. Both observations thus suggest that normalization can arise when the contribution of context effects that dominate peripheral auditory processing may be limited. With respect to the potential role of top-down modulations from regions outside of the auditory cortex, inferior frontal and sensorimotor cortex have been suggested to be involved in the resolving of perceptual ambiguities in speech perception[48,49] and could be expected to play a role in normalization too. Here we observed considerable activation in these regions, but they did not display normalization during the processing of the target sounds (see Figure S4). While tentative, these combined findings highlight the auditory cortex as the most likely locus for the emergence of normalization of speech sounds at this stage.

The current experiment involved data from cortical sites in both the left and right hemispheres. It has previously been demonstrated that the right hemisphere is more strongly involved in the processing of voice information[50,51]. Here, normalization was observed in left and right hemisphere patients (Figure 2f). Importantly, however, data included only two left hemisphere and three right hemisphere patients in total, so no strong conclusions regarding lateralization can be drawn based on this dataset.

Despite broadly observed normalization of vowel representations, responses were not completely invariant to speaker differences during the context sentences (see for example the behavior of example electrode e2 in Figure 3b which displays a preference for the Low F1 sentence though most of the context window: i.e., it is not fully normalized). And indeed, our (and previous[8,9,28]) findings show that, even for target sound processing, surrounding contexts never results in *complete* normalization. While the F1 in the context sentences differed by roughly 400 Hz, the normalization effect only induces a shift of ∼50 Hz in the position of the category boundaries (in behavior and in neural categorization). Normalization should thus be seen as a mechanism that biases processing in a context-dependent direction, but not one that fully constrains processing. Furthermore, context-based normalization is not the only means by which listeners tune-in to specific speakers. Listeners categorize sound continua differently when they are merely told they are listening to a man or a woman, demonstrating the existence of normalization mechanisms that do not rely on acoustic context[52]. In addition, formant frequencies are perceived in relation to other formants and pitch values in the current signal, because those features are correlated within speakers (e.g., people with long vocal tracts typically have lower pitch and lower formant frequencies overall). These “intrinsic” normalization mechanisms have been shown to affect auditory cortex processing of vowels as well[53–57]. Tuning-in to speakers in everyday listening is thus the result of the combination of at least these three distinct types of normalization[10].

To conclude, the results presented here reveal that the processing of vowels in auditory cortex becomes rapidly influenced by speaker-specific properties in preceding context. These findings add to recent literature that is ascribing a range of complex acoustic integration processes to the broader auditory cortex, suggesting that it participates in high-level encoding of speech sounds and auditory objects[14,25,58–60]. Recently, it has been demonstrated that patches of parabelt auditory cortex represent speaker-invariant contours of intonation that speakers use to focus on one or the other part of a sentence[61]. The current findings build on these and demonstrate the emergence of speaker-normalized representations of acoustic-phonetic features and phonemes, the most fundamental building blocks of spoken language. This context-dependence allows auditory cortex to partly resolve the between-speaker variance present in speech signals. These features of auditory cortex processing underscore its critical role in our ability to understand speech in the complex and variable situations that we are exposed to every day.

## Materials and Methods

### Patients

A total of five human participants (2 male; all right handed; mean age 30.6 years), all native Spanish speaking (the US hospital at which participants were recruited has a considerable Spanish speaking patient population), were chronically implanted with high-density (256 electrodes; 4 mm pitch) multi-electrode cortical surface arrays as part of their clinical evaluation for epilepsy surgery. Arrays were implanted subdurally on the peri-Sylvian region of the lateral left (n = 2) or right (n = 3) hemispheres. Placement was determined by clinical indications only. All participants gave their written informed consent before the surgery, and had self-reported normal hearing. The study protocol was approved by the UC San Francisco Committee on Human Research. Electrode positions for reconstruction figures were extracted from tomography (CT) scans and co-registered with the patient’s MRI.

### Stimulus synthesis

Details of the synthesis procedure for these stimuli have been reported previously[8]. All synthesis was implemented in Praat software[62]. In brief, using source-filter separation, the formant tracks of multiple recordings of clear “sufu” and “sofo” were estimated. These estimates were used to calculate a single average time-varying formant track for both words, now representing an average of the formant properties over a number of instances of both [o] and [u]. The height of only the first formant of this filter model was increased and decreased across the whole vowel to create the new formant models for the continuum from [u] to [o] covering the distance between endpoints in 6 steps. These formant tracks were combined with a model of the glottal-pulse source to synthesize the speech sound continuum. Synthesis parameters thus dictated that all steps were equal in pitch contour, amplitude contour and had identical contours for the formants higher than F1 (note that F1 and F2 values in Figure 1a and S1 reflect measurements of the resulting sounds, not synthesis parameters). The two context conditions were created through source-filter separation of a single spoken utterance of the sentence “a veces se halla” (“at times she feels rather…”). The first formant of the filter model was then increased or decreased by 100 Hz and recombined with the source model following similar procedures as for the targets.

### Procedures

The participants were asked to categorize the last words of a stimulus as either “sufu” or “sofo”. Listeners responded using the two buttons on a button box. The two options “sufu” and “sofo” were always displayed on the computer screen. Each of the 6 steps of the continuum was presented in both the low- and high-F1 sentence conditions. Context conditions were presented in separate mini-blocks of 24 trials (6 steps * 4 repetitions). Participants participated in as many blocks as they felt comfortable with.

### Data acquisition and preprocessing

Cortical Local Field Potentials (LFPs) were recorded and amplified with a multichannel amplifier optically connected to a digital signal acquisition system (Tucker-Davis Technologies) sampling at 3,052 Hz. The stimuli were presented monaurally from loudspeakers at a comfortable level. The ambient audio (recorded with a microphone aimed at the participant) along with a direct audio signal of stimulus presentation were simultaneously recorded with the ECoG signals to allow for precise alignment and later inspection of the experimental situation. Line noise (60Hz and harmonics at 120 and 180 Hz) was removed from the ECoG signals with notch filters. Each time series was visually inspected for excessive noise, and trials and or channels with excessive noise or epileptiform activity were removed from further analysis. The remaining time series were common-average referenced across rows of the 16×16 electrode grid. The time-varying analytic amplitude was extracted from eight bandpass filters (Gaussian, with logarithmically increasing center frequencies between 70–150 Hz, and semi-logarithmically increasing bandwidths) with the Hilbert transform. High-gamma power was calculated by averaging the analytic amplitude across these eight bands. The signal was subsequently down-sampled to 100Hz. The signal was z-scored based on the mean and standard deviation of a baseline period (from -50 to 0 ms before the onset of the context sentence) on a trial by trial basis. In the main text, high-γ will refer to this measure.

### Single-electrode encoding analysis

We used ordinary least-squares linear regression to predict neural activity (high-γ) from our stimulus conditions (target F1 steps, coded as -2.5, -1.5, -0.5, 0.5, 1.5, 2.5; and context F1, coded as -1 and 1; as well as their interaction). These factors were used as numerical predictors to neural activity that was averaged across the target window (from 70 to 570 ms after target vowel onset) or across the context window (from 250 to 1450 ms after context sentence onset –a later onset was chose to reduce the influence of large and non-selective onset responses present in some electrodes-). For each model R-squared (Rsq) provides a measure of the proportion of variance in neural activity that is explained by the complete model. The p-value associated with the omnibus F-statistic provides a measure of significance. We set the significance threshold at alpha = 0.05 and corrected for multiple comparisons using the Bonferroni method, taking individual electrodes as independent samples. Figure S2a & b demonstrate that most of the variance in the context was explained by the factor context F1. During the target window however, both target F1 and context F1 explain a considerable portion of the variance. The interaction term was included to accommodate a situation where the context effect is more strongly expressed on one side of the target continuum than the other (see e.g., figure 2b, where the context effect is larger towards “sofo”), but is not further interpreted here.

For Figures 2d and Figure 3c, linear correlations between signed t-statistics of target F1 preferences and context effects (Figure 2d) or context preferences (Figure 3c) were computed over all significant (9 [corrected] + 28 [uncorrected] = 37) electrodes. For Figure S6a, linear correlations were computed separately for those electrodes that displayed significant selectivity for Target F1 and Context F1 (n = 9; (marked with a black-and-white outline in S6a; *r* = -0.73; *p* = 0.03), and for electrodes that displayed selectivity to Target F1 only (n = 28; S6a; *r* = -0.61; *p* = 5.07*10^-4^). For Figure S6b the same approach was applied for each high-gamma sample separately.

### Cluster-based permutation analyses

For single example electrodes, a cluster-based permutations approach was used to assess significance of differences between two event related high gamma time courses (Figure 2c and Figure 3b; following the method described in[63]). For each permutation, labels of individual trials were randomly assigned to data (high-y time courses), and a t-test was performed for each timepoint. Next, for each time point (across all 1000 permutations) a criterion value was established (the highest 95% of the [absolute] t-values for that timepoint). Then, for each permutation, it was established when its value reached above the criterion value and for how many samples it remained above criterion. A set of subsequent timepoints above criterion is defined as a cluster. Then, for each cluster the t-values were summed, and this value was assigned to that entire cluster. For each permutation only the largest (i.e., highest summed cluster value) was stored as a single value. This resulted in a distribution of maximally 1000 cluster values (some permutations may not result in any significant cluster and have a summed t-value of 0). Then, using the same procedure, the size of all potential clusters was established for the real data (correct assignment of labels), and it was established whether the size of each cluster was larger than 95% of the permutation-based cluster values. p < 0.001 indicates that the observed cluster was larger than all permutation based clusters.

### Stimulus classification

Linear Discriminant Analysis (LDA) models were trained to predict the stimulus from the neural population responses evoked by the stimuli. Per participant a single model was trained on all endpoint data, which was then used to predict labels for the ambiguous items. To predict stimulus class for the endpoint stimuli (steps 1 and 6) a leave-one-out cross validation procedure was used to prevent overfitting. Model features (predictors) consisted of the selected timepoint*electrode combinations per participant.

For the analyses (Figures 2; Figure S3; Figure S4) training data consisted of high-y data averaged across a 500ms time window starting 70ms after target vowel onset (the target vowel was the first point of acoustic divergence between targets).

In the analyses, all task-related electrodes for a given participant (and region-of-interest, see Figure S4) were selected. Trial numbers per participant are listed in Table S1. The analysis displayed in Figure 2 and Figures S3 and S4 hence relied on a large number of predictors (electrodes * timepoints). While a large amount of predictors could result in overfitting, these parameters led to the highest proportion of correct classification for the endpoints (76% correct, see Figure S1b). High endpoint classification performance is important to establish the presence of normalization, but does not affect the extent of observed normalization, because the normalization effect is orthogonal. Importantly, in all analyses classification scores were only obtained from held-out data, preventing the fitting of idiosyncratic models. In addition, averaging across time (hence decreasing the number of predictors) led to qualitatively similar (and significant) effects for the important comparisons reported in this paper. Classification analyses resulted in a predicted class for each trial. These data were used as input for a generalized logistic linear mixed effects model.

### Generalized Linear Mixed effects regression of classification data (glmer)

For the analyses that assessed the effects of target stimulus F1 and context F1 on proportion of “sofo” responses (both behavioral and neural-classifier-based), the models had Target F1 (contrast coded, with the levels -2.5; -1.5; -0.5; 0.5; 1.5; 2.5) and Context F1 (levels -1; 1) entered as fixed effects, and uncorrelated by-patient slopes and intercepts for these factors as random effects.

For the analysis of the behavioral data, we observed more *sofo* responses towards the sofo end of the stimulus continuum (B_Target F1_ = 1.89, z = 3.62, p < 0.001). Moreover, we observed an effect of context as items along the continuum were more often perceived as *sofo* (the vowel category corresponding to higher F1 values) after a low F1 voice (Speaker A) than after a high F1 voice (Speaker B) (B_Context F1_ = -1.71, *z* = -3.15, p = 0.002).

For the analyses of neural representations the dependent variable consisted of the classes predicted by LDA stimulus classification described above. For the overall analysis including temporal lobe electrodes, the model revealed significant classification of the continuum (B_Target F1_ = 0.50, *z* = 13.18, p < 0.001), suggesting reliable neural differences between the endpoints. Furthermore, an effect was also found for the factor Context on the proportion of “sofo” classifications (B_Context F1_ = -0.28, *z* = -4.67, p < 0.001), reflecting the normalization effect of most interest. For the analysis focusing on the dorsal and frontal electrodes a significant effect of Step was observed, that is, significant classification of the continuum (B_Target F1_ =0.20, *z* = 6.04, p< 0.001), but no significant influence of context (B_Context F1_= 0.02, *z* = 0.31 p = 0.76) see S4C for further detail.

### Multidimensional scaling

The neural dissimilarity between all 12 pairs of target items was measured by computing leave-one-out LDA classification scores between each target pair. Here, a high classification score reflects different neural representations, a low score reflects similarity. The resulting 12*12 dissimilarity matrices were averaged across participants (Figure S5a). The across-participant average of the classification-based distance matrices was projected in Multidimensional Scaling space. The first (i.e., main) dimension reflected stimulus step (see Figure S5b), indicating that this is indeed the most dominant property of the selected electrode population. Importantly, this dimension also reflected normalization

## Author Contributions

M.J.S. and K.J., conceived the study. M.J.S. designed the experiments, generated the stimuli and analyzed the data. M.J.S., N.F. and E.F.C. collected the data. M.J.S. and N.F. interpreted the data and wrote the manuscript. M.J.S., N.F., K.J., and E.F.C. edited the manuscript.

## Code availability

These results were generated using code written in Matlab. Code will be publicly available through the Open Science Framework osf.io/u7xnp.

## Data availability

The data that support the findings of this study will be publicly available through the Open Science Framework osf.io/u7xnp.

## Acknowledgements

We are grateful to Matthew Leonard for commenting on an earlier version of this manuscript. This work was supported by European Commission grant FP7-623072 (MJS); and NIH grants F32-DC013486, R00-NS065120, DP2-OD00862, and R01-DC012379 (E.F.C.); and F32DC015966 (N.P.F.).

## Supporting information

**Figure S1:**
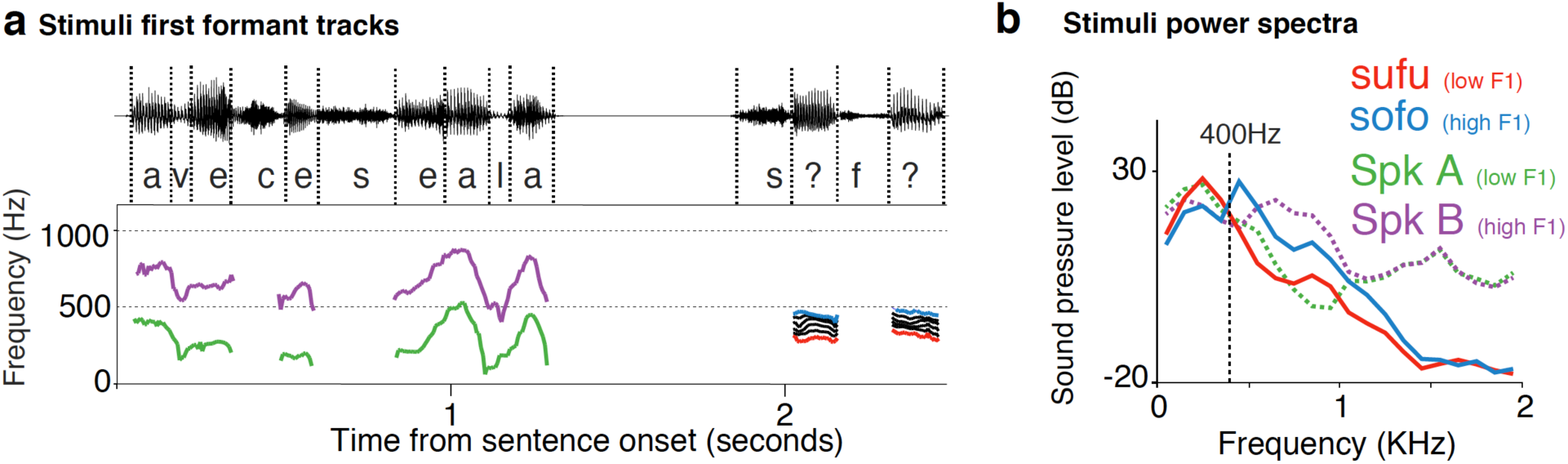
Contexts and targets display similar spectral differences. **a**) First formant tracks for the voiced portions of the synthesized materials. In the annotation “?” indicates one of the target vowel steps. **b)** A similar spectral relation exists between the two endpoint targets /u/ vs. /o/ and the context sentences. /u/ and the low F1 speaker have more dominant low frequency components (i.e., below 400 Hz) in the spectrum than /o/ and the High F1 speaker.

**Figure S2:**
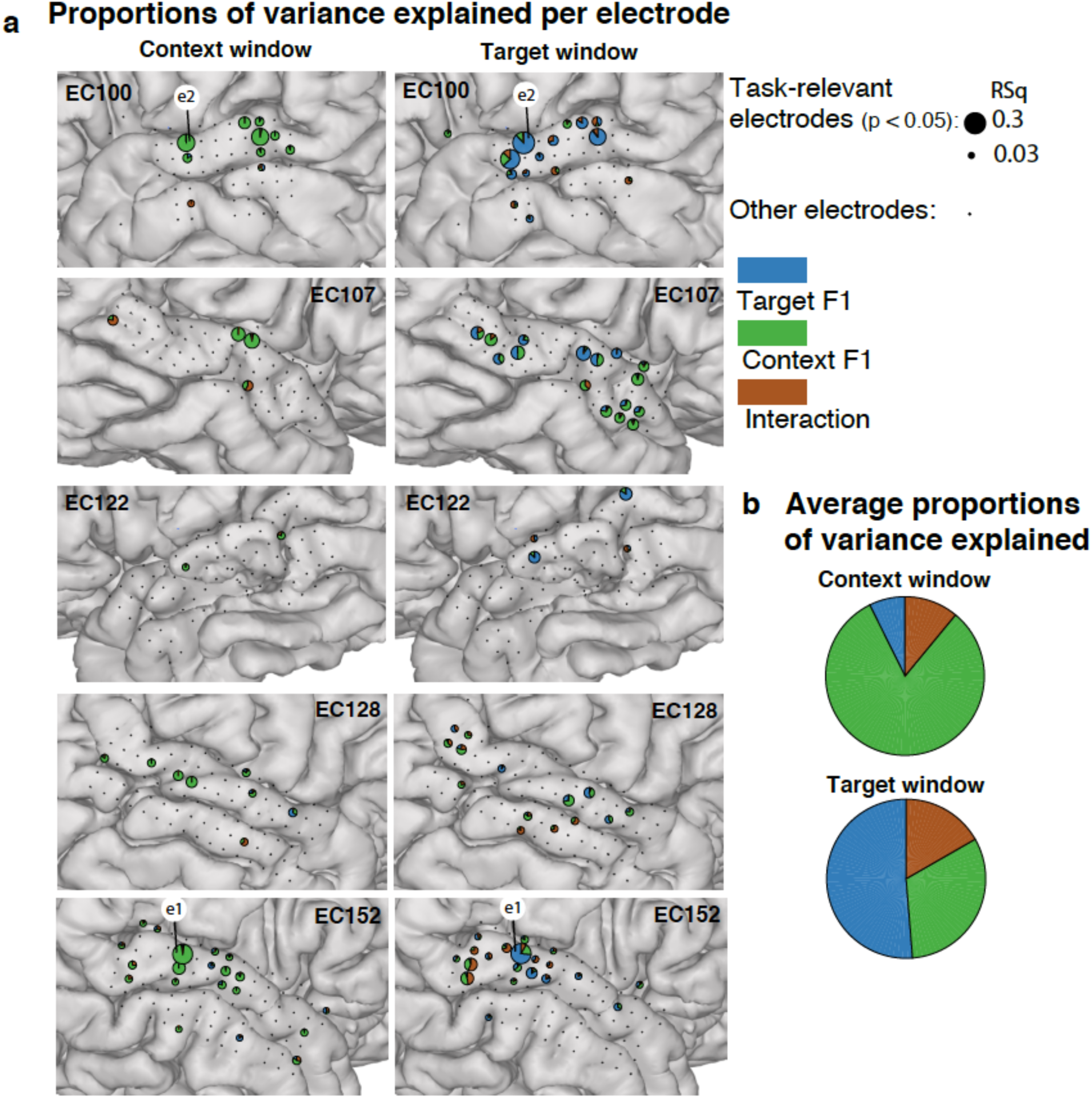
Context F1 influences cortical activity during target processing. a) Map of context and target encoding for all task-related electrodes of all subjects (temporal lobe only), both during the context window (left column) and during the target window (right column). The areas of the pie-charts are proportional to the total variance explained. Wedges show the relative variance explained by each factor (stimulus dimension) for each significant electrode. **b**) Weighted average proportion of variance explained by main effects and interactions across all significant electrodes (across all 5 patients). During the context window, context properties explain the large majority of variance. During the target window, the context stimulus properties still explain a considerable portion of the variance.

**Figure S3:**
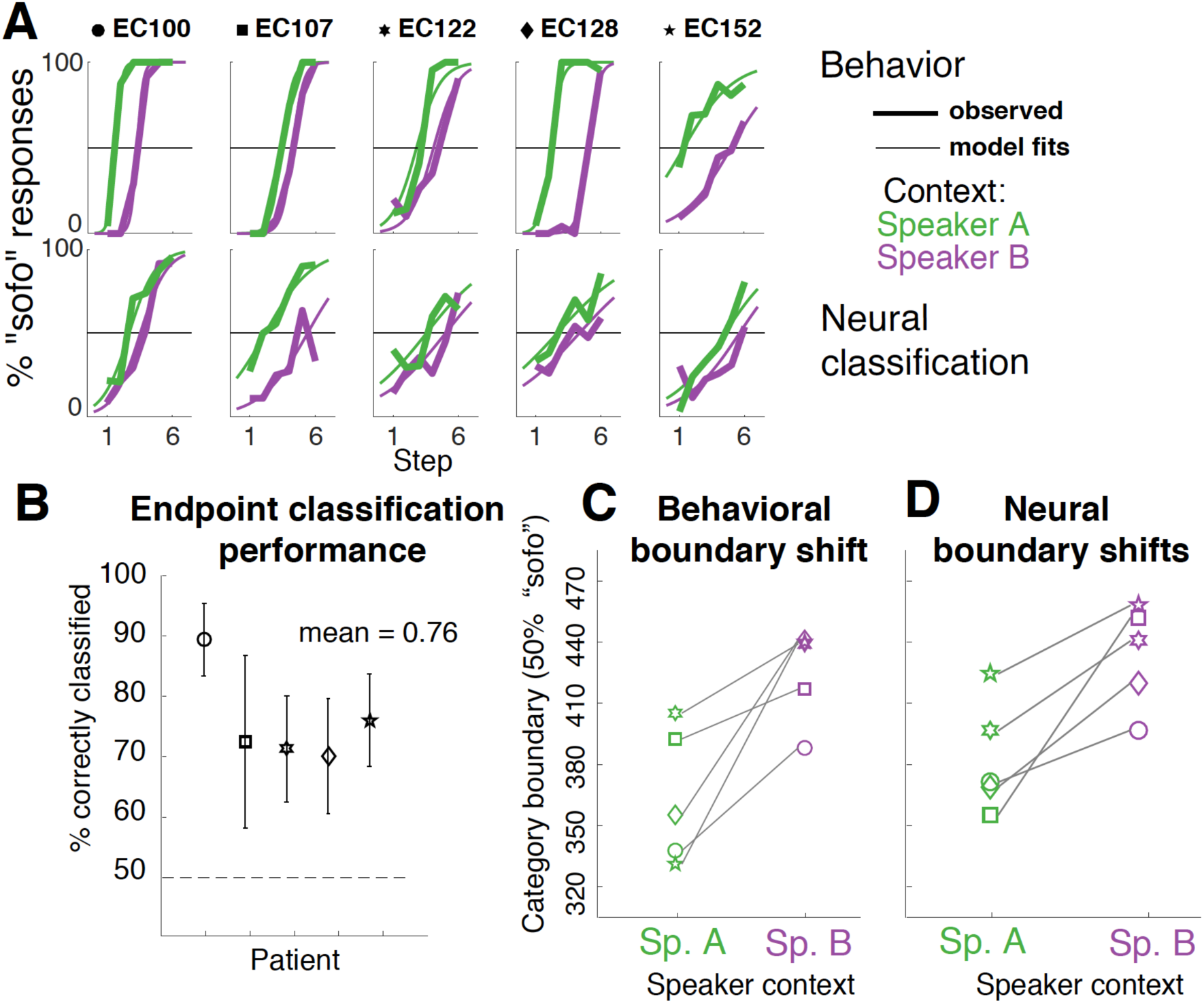
All participants display context effects in both their behavior and neural responses. **a**) Top row: Observed mean proportions of “sofo” responses across the steps of the continuum in both context conditions for all participants (thick lines). Model fits (thin lines) are used to estimate the 50% category boundary per condition per participant (used for panel c). Bottom: same as in top panel but for the neural classification data (thick lines reflect LDA-based predictions). **b**) overall percentage correct (leave-one-out) classification on the endpoints per participant (with bootstrapped 95% CI). **c**) By-participant indications of the estimated 50% category boundaries in the two context conditions based on behavior. **d**) same as c but for 50% category boundary estimates of the neurally-based classification (i.e., identical to Figure 2d, but reproduced for comparison to Fig. S3c).

**Figure S4:**
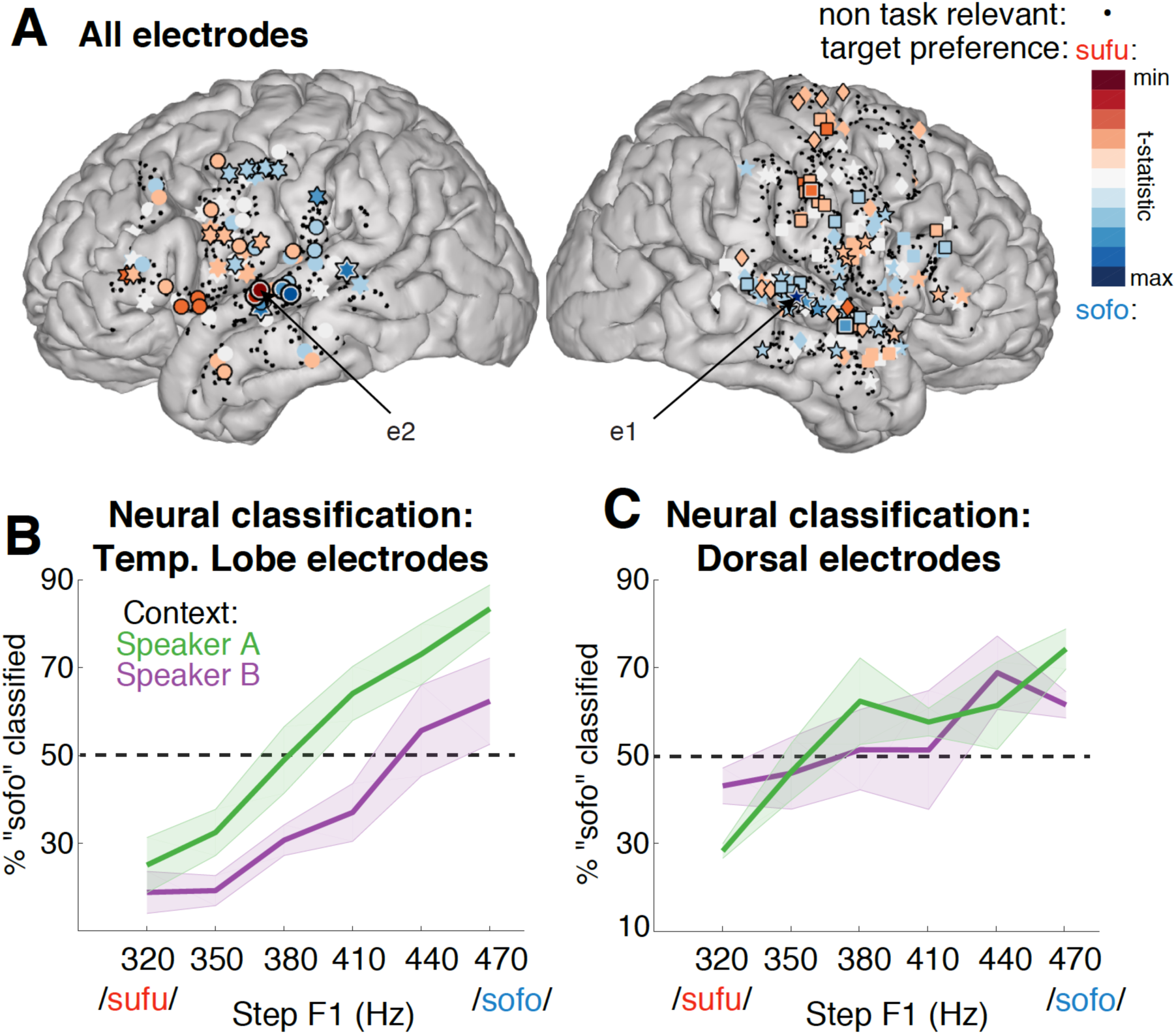
Normalization is observed only in temporal lobe regions. a) Target vowel selective electrodes are congregated on the temporal lobe, although some are also found in frontal regions. **b**) when including temporal electrodes, LDA classification results (averaged across participants) reveal a strong context effect. **c**) no context effect is observed for LDA classification results based on electrodes from dorsal (i.e., all non-temporal) regions.

**Figure S5:**
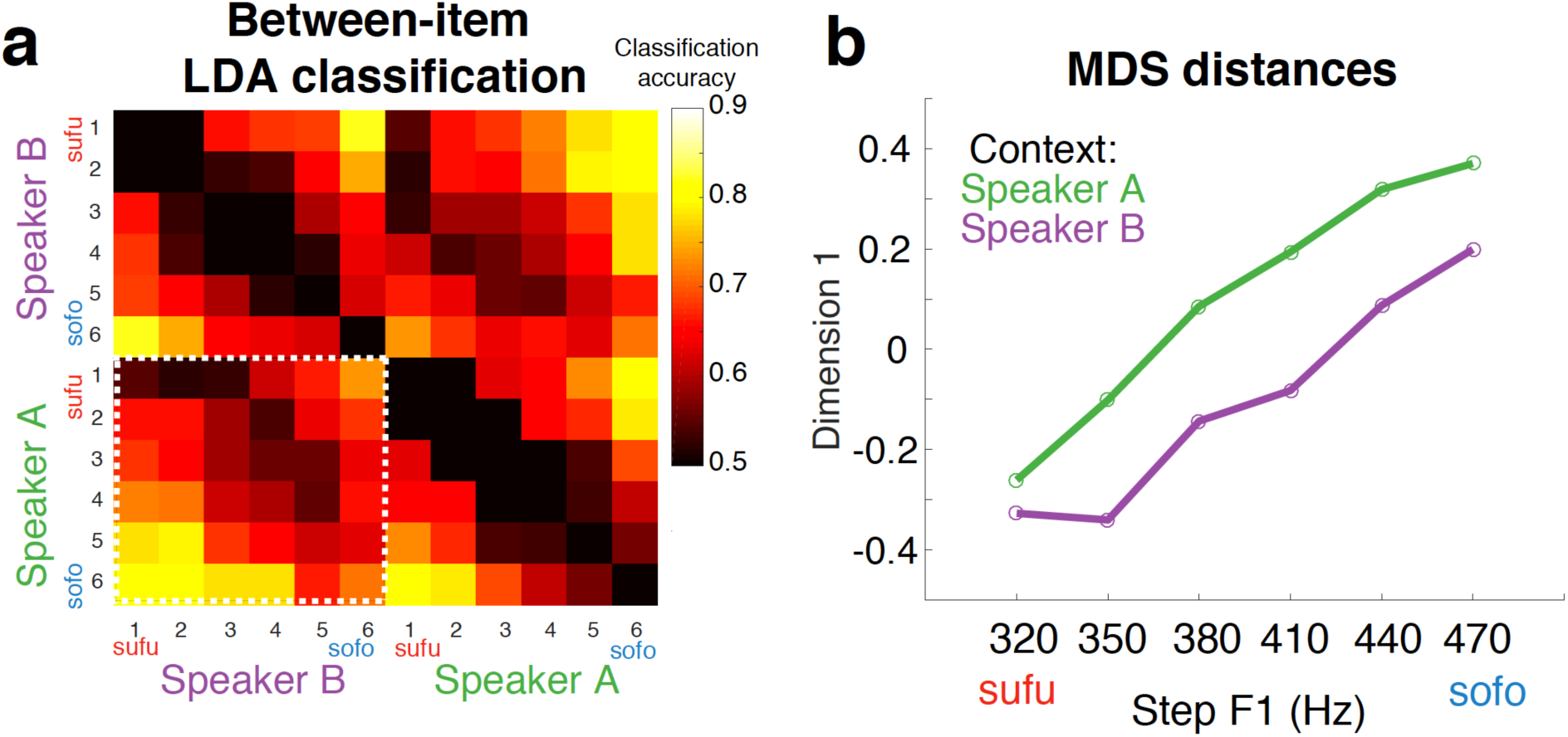
Between stimulus neural distances reflect normalization. a)Distance matrix based on between-stimulus leave-one-out LDA classification of a two-class classifier. High classification performance reflects large neural distances. The lower left corner (involving the between-context comparisons; marked with a white-dashed outline) demonstrates normalization: e.g., step 2 of Speaker A is most similar to step 4 of Speaker B, etc. Under veridical processing, smallest distances would follow the sub-diagonal (with the smallest distances for the comparisons of 1 vs. 1; 2 vs. 2; etc.) **b)** Multidimensional scaling (MDS) based on the distance matrix (in a) reveals that target stimulus F1 is the main factor (D1) determining neural dissimilarity. Critically, this dimension is also influenced by context F1. That is, it reflects normalization.

**Figure S6:**
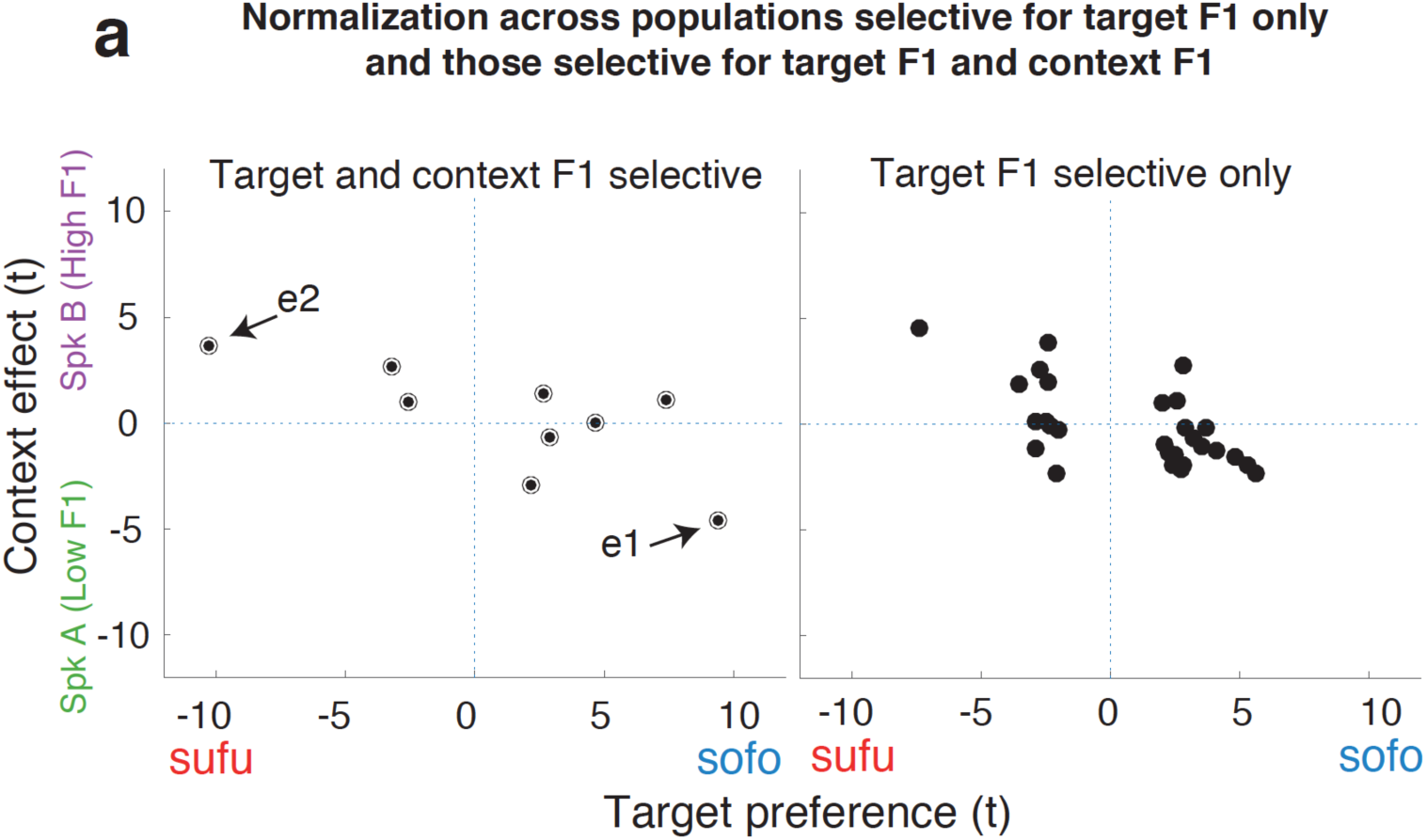
Normalization across target F1 selective population types. Panels displays the same relation as Figure 2d, (the relation between target preference and context effect; i.e., normalization), but for two types of populations. Normalization is observed for electrodes that are most clearly selective for acoustic-phonetic features (instead of phonemes) since they display preferences for both target F1 and context F1 (left panel; black-and-white outline; target and context materials contain different phonemes). For completeness; normalization is also observed for those electrodes that display target F1 preferences only (right panel; black fill). See Figure 3 for reference.

## SUPPLEMENTARY TABLES

**table S1:** Trial counts

**S1: Trial counts**
| 769 | Item | EC100 | EC107 | EC122 | EC128 | EC152 |
| --- | --- | --- | --- | --- | --- | --- |
| 770 | High_1 | 24 | 9 | 28 | 23 | 33 |
| 771 | High_2 | 24 | 9 | 26 | 23 | 26 |
| 772 | High_3 | 24 | 12 | 31 | 24 | 31 |
| 773 | High_4 | 23 | 11 | 27 | 24 | 27 |
| 774 | High_5 | 24 | 11 | 29 | 19 | 29 |
| 775 | High_6 | 24 | 9 | 27 | 22 | 26 |
| 776 | Low_1 | 23 | 11 | 25 | 21 | 31 |
| 777 | Low_2 | 24 | 12 | 27 | 21 | 29 |
| 778 | Low_3 | 24 | 9 | 23 | 20 | 33 |
| 779 | Low_4 | 23 | 8 | 30 | 20 | 31 |
| 780 | Low_5 | 24 | 10 | 25 | 19 | 31 |
| 781 | Low_6 | 23 | 11 | 28 | 21 | 31 |

**table S2:** Electrode type counts (temporal lobe)

|  |  |  |  |  |
| --- | --- | --- | --- | --- |
| 787 | EC100 | 21 | 10(4) | 8(3) |
| 788 | EC107 | 19 | 7(1) | 4(2) |
| 789 | EC122 | 6 | 3(2) | 3(0) |
| 790 | EC128 | 20 | 6(0) | 10(2) |
| 791 | EC152 | 32 | 11(2) | 16(4) |

